# Endogenous synthesis of pyrethrins by cannabis

**DOI:** 10.1101/169417

**Authors:** Adrian Devitt-Lee, Douglas R. Smith, David Chen, Kevin McKernan, Simone Groves, Ciaran McCarthy

## Abstract

Pyrethrins are a class of natural terpenoid pesticides produced by *Tanacetum cinerariifolium*, commonly known as chrysanthemum. Here we present evidence that cannabis may be able to produce pyrethrins endogenously. Flower from a cannabis plant grown in a closed hydroponic environment contained 2.48 parts per million pyrethrin I by weight. A comparison of the genetics of *T. cinerariifolium* and *Cannabis* demonstrates Cannabis homologues of the genes that contribute to pyrethrins production in *T. cinerariifolium*. This provides a plausible pathway for the biosynthesis of pyrethrins in cannabis. Although preliminary, these data indicate a potentially significant confounding variable in both cannabis research and regulations on allowable pyrethrins residues in cannabis products.

## Introduction

Pyrethrins are a group of natural insecticides produced by *Tanacetum cinerariifolium*. Pyrethrins consist of six related compounds -- pyrethrin I/II, jasmolin I/II, and cinerin I/II--formed from the esterification of one of three rethrolone moieties and either chrysanthemic or pyrethric acid (see figure 1). Pyrethrin I is often used as the model for pyrethrins and synthetic derivatives of pyrethrins, called pyrethroids.

**Figure 1.**
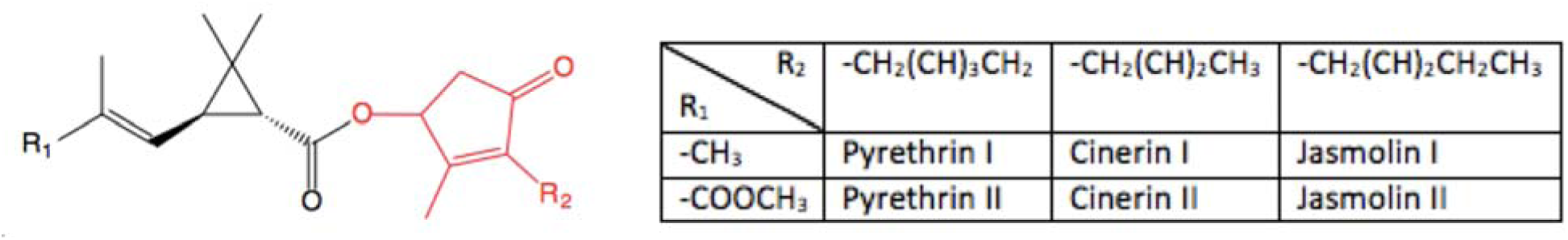
Structure of pyrethrins. The acid derivative is highlighted in red and the rethrolone in black. R_1_ is a methyl group (chrysanthemic acid) or methyl ester (pyrethric acid), and R_2_ is 2-butene, 2-pentene, or 2,4-pentdiene. The corresponding compound names are given on the right.

Biologically, pyrethrins are formed from the convergence of the oxylipin and a monoterpene pathway, two pathways that are highly conserved among plants. Pyrethrin I’s biosynthesis is shown in figure 2.^1-2^ To form the acid monoterpene, an isoprene (IPP) unit is first isomerized to dimethyl allyl pyrophosphate (DMAPP). Chrysanthemyl diphosphate synthase (CDS) combines two DMAPP units to form chrysanthemyl diphosphate, followed by a series of oxidations converting this terpene to chrysanthemic acid.^3^ Finally, chrysanthemic acid is conjugated to coenzyme A and transferred to a rethrolone by *T. cinerariifolium* GDSL lipase (Tc GDSL lipase; TcGLIP). The rethrolones are formed from the oxylipin pathway, beginning with the oxidation of linolenic acid. The oxylipin pathway appears to occur in both the leaves and flower of *T. cinerariifolium*, whereas chrysanthemic acid-synthesizing enzymes predominate in the glandular trichomes of the flower.^1^

The formation of the cyclopropane ring in chrysanthemic acid by CDS, as well as the final esterification by TcGLIP are two key steps in the production of pyrethrins. CDS appears to be expressed in the flowers of chrysanthemum but not other parts of the plant, allowing the production of pyrethrins to be localized to trichomes.^4^ There is some evidence that certain enzymes involved in the production of rethrolones are also expressed primarily in glandular trichomes.^5^ TcGLIP is generally expressed in the pericarp of seeds rather than trichomes, but its expression in trichomes can be induced.^4,6^ Relative expression of these genes depends on the plant stressors present, as pyrethrin production is a defense mechanism. The genetics of many of the enzymes involved in producing pyrethrins, including CDS and TcGLIP have been described previously.^1,6-9^

## Methods

### Growing cannabis

Mature CBD Medihaze seedlings were grown in rockwool cups adjacent a 20 gallon hydroponic tank at sea level from a seed from CBD Crew. The plant was grown with a Spectrum King 400 plus LED with a 16:8 vegetative light cycle and a 12:12 flower cycle. A tower garden closed loop aeroponic system was used to deliver eight 15-minute waterings per day. 400ml of Tower Garden Mineral Blend A and B nutrients were used to initially equilibrate the E.C under 3 and pH under 6.5. These nutrients were complimented with 280ml of General Hydroponics FloraNova Bloom (4-8-7) after 5 weeks of vegetative growth.

### LC-MS-MS

Cannabis flower was homogenized and 531 mg of the homogenate was dissolved in 10 mL of methanol with 0.1% formic acid. This solution was diluted by a factor of 20 in acetonitrile and triphenyl phosphate, then injected into the LC. The MS is run using Analyst and MultiQuant softwares, both available through AB Sciex. Based on the calibration curve for this experiment, the concentration of pyrethrins in flower is [Pyrethrin I] = (*A*∗2.86∗10^−2^ + 132)/*m*, where the concentration is given in μg/g, *m* is the sample mass in mg, and *A* is the peak area for the m/z = 329.3 to 161.1 transition.

### Genetic analysis

The NCBI non-redundant protein sequence database (nr database) was searched for enzyme names that matched those in the *Tanacetum cinerariifolium* pyrethrin biosynthetic pathway described by Sakamori et al.^2^ The resulting sequences (from a variety of different organisms) were used as queries for BLAT searches of the canSat3 assembly using the tool available at the Cannabis Genome Browser Gateway (http://genome.ccbr.utoronto.ca/cgi-bin/hgGateway).^10-11^ The set of scaffolds representing the resulting matches were examined to identify probable coding transcripts. Transcript sequences were downloaded and translated to identify the encoded protein sequence. The resulting sequences were used to search the nr database, using BLASTP, to identify best hits and verify the consistency of enzyme names across different plant species. A summary of the results of these searches is provided in Table 1 and Supplementary Table 1.

## Results

Cannabis grown hydroponically in a closed system was tested for pyrethrin I using LC-MS-MS. The peaks for the pyrethrin standard and cannabis flower are shown in figure 3. No pesticides were detected in the water (data not shown), but a significant peak corresponding to pyrethrin I was observed in cannabis flower, indicating 2.48 μg pyrethrin I/g flower. This was quantified using the transition from m/z = 329.3 to 161.1. The former mass is the protonated pyrethrin I, whereas the latter is a pyrethrolone ion lacking its hydroxyl group. A slight peak was detected in cannabis leaves suggesting the presence of pyrethrins, but this was not quantified due to the noisiness of the sample (data not shown). This is unsurprising, given that pyrethrins are produced at 10-20 fold lower concentrations in the leaves of *T. cinerariifolium*.^4^ The humps visible in figures 3A/C/D are likely due to the coelution of pyrethrins.

### Identification of putative pyrethrin biosynthetic enzyme coding sequences in *Cannabis*

Next we identified a potential biological mechanism for the production of pyrethrins by cannabis. A summary of the comparison between *T. cinerariifolium* and *Cannabis* transcripts are provided in Table 1 and Supplementary Table 1. The translated sequences of *Cannabis* transcripts PK18899.1 and PK09506.1 were found to be truncated at their N-termini by 46 aa and 174 aa, respectively, relative to their homologs.

**Table 1.**
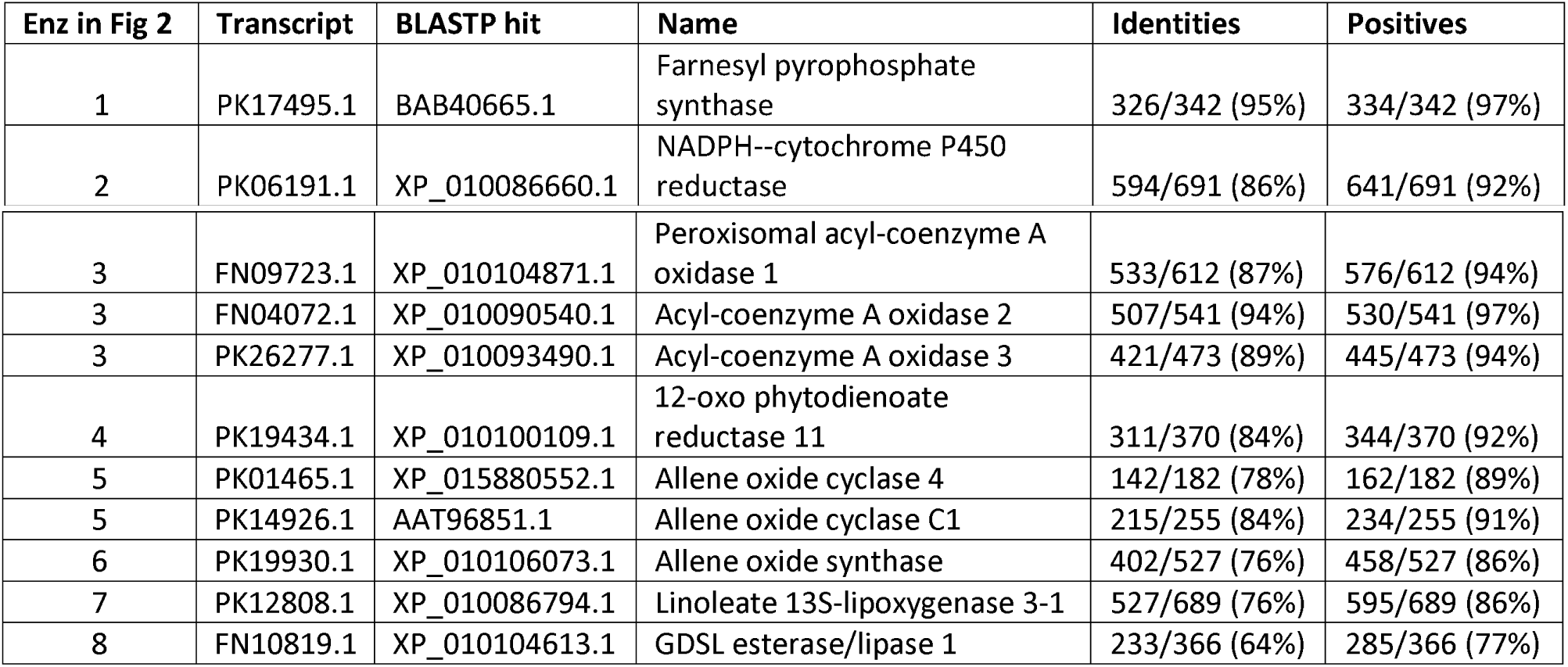
BLASTP results of *Cannabis* transcripts against Genbank nr protein database.

| Enz in Fig 2 | Transcript | BLASTP hit | Name | Identities | Positives |
| --- | --- | --- | --- | --- | --- |
| 1 | PK17495.1 | BAB40665.1 | Farnesyl pyrophosphate synthase | 326/342 (95%) | 334/342 (97%) |
| 2 | PK06191.1 | XP_010086660.1 | NADPH--cytochrome P450 reductase | 594/691 (86%) | 641/691 (92%) |
| 3 | FN09723.1 | XP_010104871.1 | Peroxisomal acyl-coenzyme A oxidase 1 | 533/612 (87%) | 576/612 (94%) |
| 3 | FN04072.1 | XP_010090540.1 | Acyl-coenzyme A oxidase 2 | 507/541 (94%) | 530/541 (97%) |
| 3 | PK26277.1 | XP_010093490.1 | Acyl-coenzyme A oxidase 3 | 421/473 (89%) | 445/473 (94%) |
| 4 | PK19434.1 | XP_010100109.1 | 12-oxo phytodienoate reductase 11 | 311/370 (84%) | 344/370 (92%) |
| 5 | PK01465.1 | XP_015880552.1 | Allene oxide cyclase 4 | 142/182 (78%) | 162/182 (89%) |
| 5 | PK14926.1 | AAT96851.1 | Allene oxide cyclase C1 | 215/255 (84%) | 234/255 (91%) |
| 6 | PK19930.1 | XP_010106073.1 | Allene oxide synthase | 402/527 (76%) | 458/527 (86%) |
| 7 | PK12808.1 | XP_010086794.1 | Linoleate 13S-lipoxygenase 3-1 | 527/689 (76%) | 595/689 (86%) |
| 8 | FN10819.1 | XP_010104613.1 | GDSL esterase/lipase 1 | 233/366 (64%) | 285/366 (77%) |

Of the *T. cinerariifolium* enzymes shown in figure 2, homologs of all but two could be identified in *Cannabis* with similarities ranging from 77-98% (Table 1). Sequences corresponding to chrysanthemic acid coA ligase could not be identified in the nr database. Nor did BLAT searches of canSat3 assembly with chrysanthemic diphosphate synthase (CDS) sequences from various organisms result in any matches. However, matches were found using closely related farnesyl diphosphate synthase (FDS) sequences, from which CDS is thought to be derived evolutionarily by duplication.^8^ A multialignment of this *Cannabis* FDS sequence with several CDS sequences shows a high degree of relatedness, but the *Cannabis* enzyme lacks a 55 amino acid extension at the N-terminus (figure 4). Multiple DDXXD motifs in FDS and CDS coordinate the DMAPP and IPP units in the active sites of these enzymes, and these are boxed in figure 4.^8^ Sites where one aspartate residue is mutated to asparagine (NDXXD or DNXXD) – which appears to be significant to the function of CDS – are also indicated.^9^ The *Cannabis* FDS gene contains a DNXXD sequence 20 amino acids upstream of the NDXXD motif in the CDS genes.

**Figure 2.**
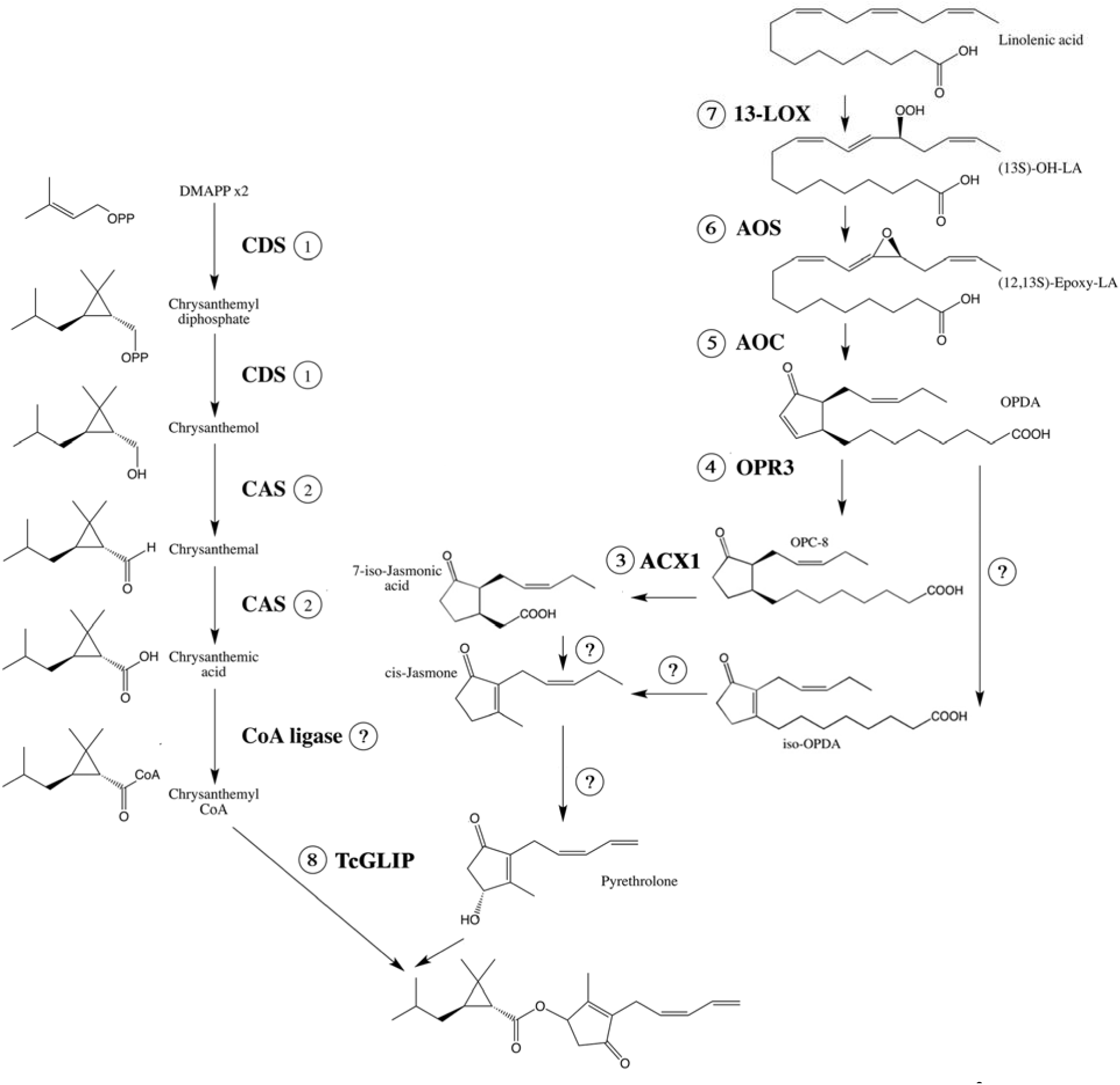
Synthesis of pyrethrin I in *Tanacetum cinerariifolium* identified by Sakamori et al.^2^ Compound names are beside their structures. Enzyme names, if known, are shown in bold alongside each arrow. Abbreviations: LA, linolenic acid; OPC-8, 3-oxo-2-(2-pentenyl)-cyclopentane-l-octanoic acid; OPDA, 12-oxo-phytodienoic acid. **ACX1**, acyl coenzyme A oxidase 1; **AOC**, allene oxide cyclase; **AOS**, allene oxide synthase; **CAS**, chrysanthemic acid synthase; **CoA ligase**, chrysanthemic acid-coenzyme A ligase; **CDS**, chrysanthemyl diphosphate synthase; **HPL**, hydroperoxide lyase; **13-LOX**, 13-lipoxygenase; **OPR3**, 3-oxo-2-(2-pentenyl)-cyclopentane-l-octanoic acid reductase 3; **TcGLIP**, Tanacetum cinerariifolium GDSL lipase.

**Figure 3.**
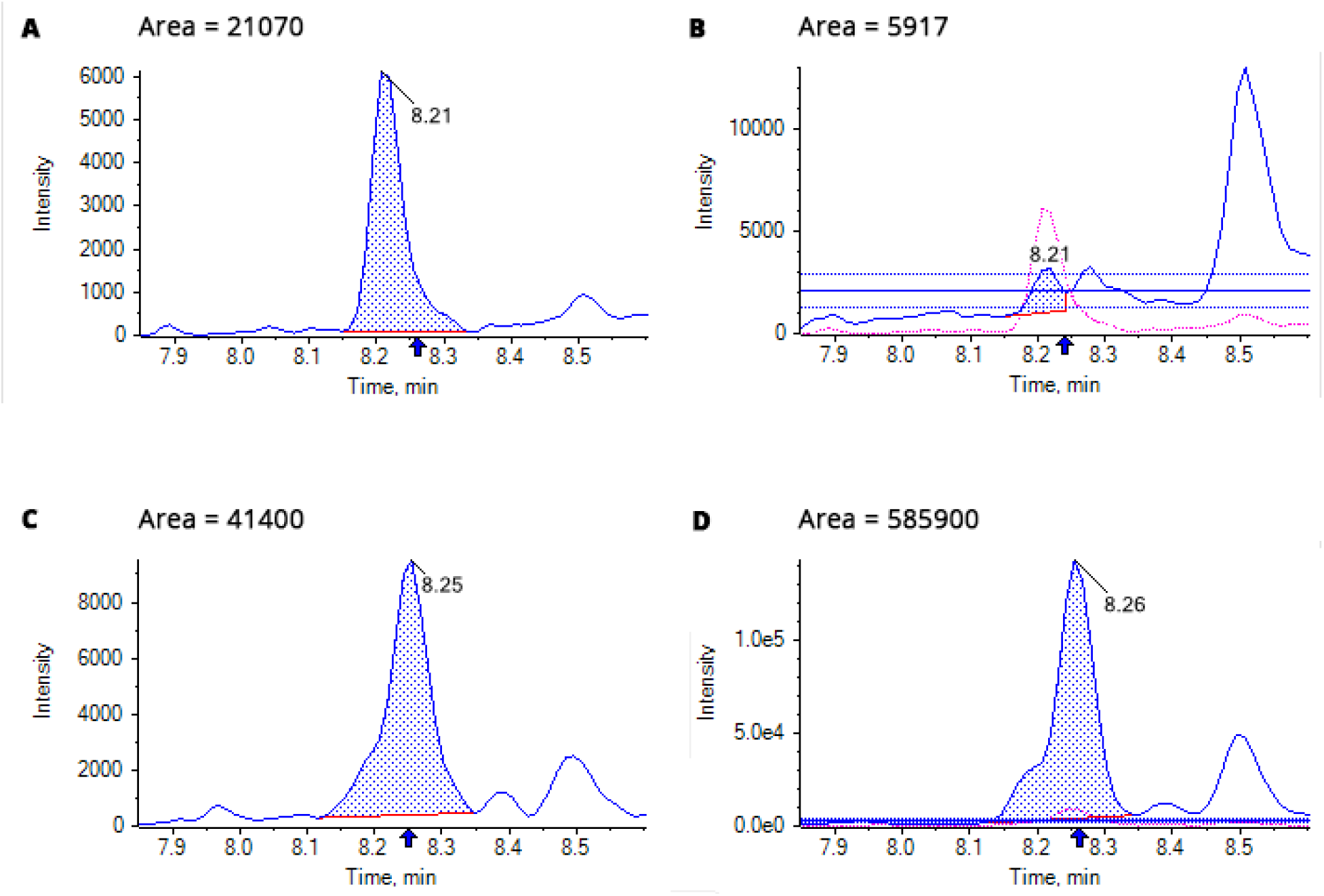
Chromatograms for the 3 ng/ml pyrethrin standard and homogenized cannabis flower. *A*: Pyrethrin standard with transitions m/z = 329.3 and m/z = 161.1. *B*: Pyrethrin standard with transitions m/z = 329.3 and m/z = 133.2. *C*: Cannabis flower with transitions m/z = 329.3 and m/z = 161.1. *D*: Cannabis flower with transitions m/z = 329.3 and m/z = 133.2.

**Figure 4.**
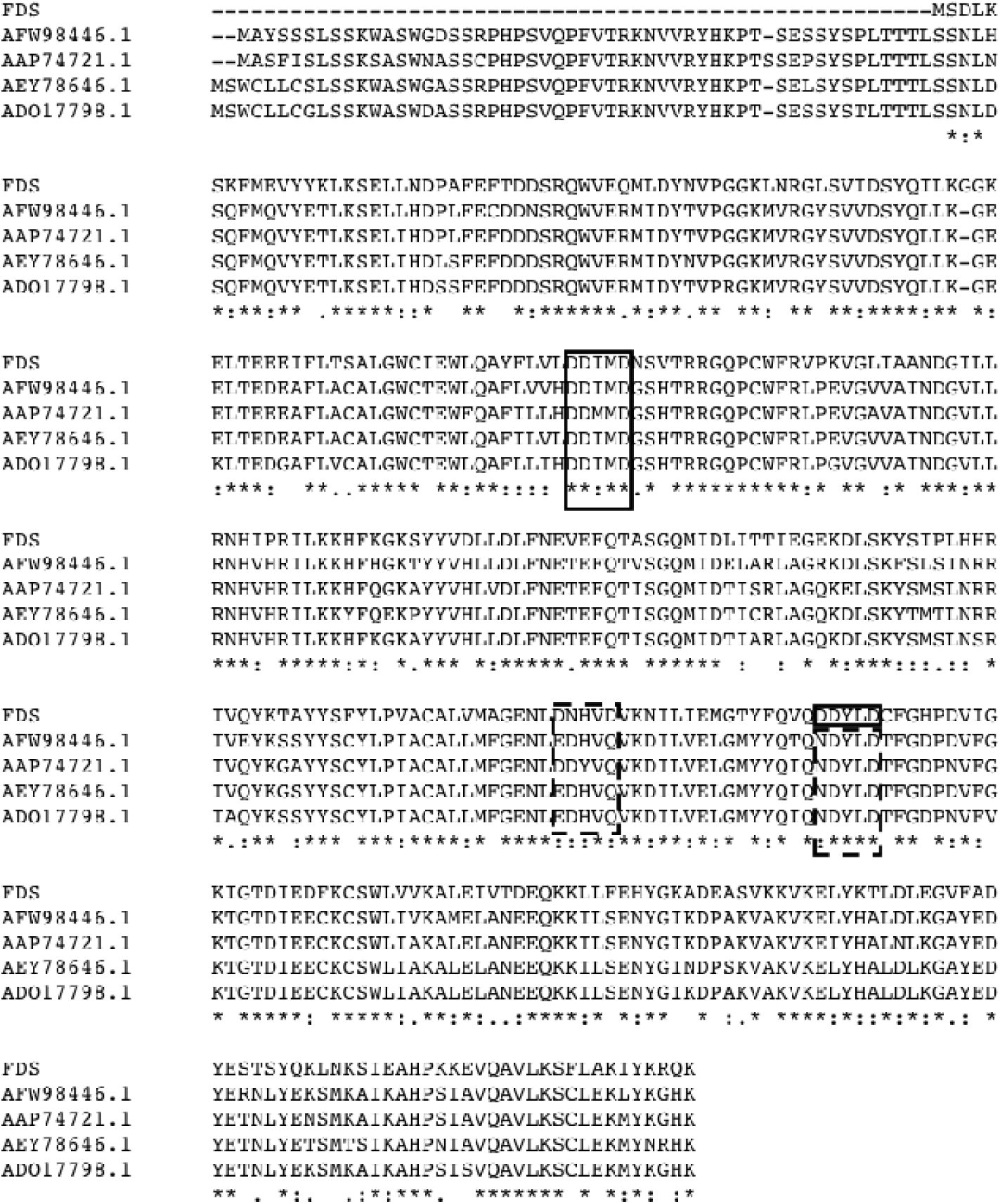
Clustal multialignment of *Cannabis* FDS to CDS sequences from *Achillea asiatica* (AFW98446.1), *Artemisia spiciformis* (AAP74721.1), *Tagetes erecta* (AEY78646.1) and *Tanacetum cinerariifolium* (ADO17798.1). DDXXD motif are boxed. DNXXD and NDXXD motifs are in dashed boxes.

Two of the transcripts identified (PK18899.1 for ACX1, and PK09506.1 for ACX2) appeared to be truncated at their 5’ ends compared to other plant sequences. The corresponding transcripts from the Finola assembly (FN09723.1 for ACX1, and FN04072.1 for ACX2) were also truncated, but by fewer amino acids, so those were selected for analysis. Finola is a common hemp variety.

The *Cannabis* homologs identified support the possible presence of the two pathways involved in pyrethrin I biosynthesis: from DMAPP to chrysanthemic acid (Chr-COOH), and from linolenic acid to 7-iso-jasmonic acid (7-IJA). The lack of published sequence data on the final enzymes in those two pathways, leading from Chr-COOH to chrysanthemoyl CoA (or from chrysanthemol pyrophosphate as described by others), and from 7-IJA to pyrethrolone, precluded us from identifying homologs of those enzymes.^12^

The last step in the pyrethrin biosynthetic pathway in *T. cinerariifolium*, condensation of chrysanthemoyl CoA and pyrethrolone, is reported to be catalyzed by a GDSL esterase/lipase, termed TcGLIP.^6,13^ Transcript PK07938.2 for GDSL lipase was fragmentary, so an alternative transcript from the Finola assembly was selected (FN10819.1). An alignment of the putative *Cannabis* GDSL lipase with the *T. cinerariifolium* GDSL lipase is shown in figure 5. Several *Cannabis* GDSL esterase/lipase homologs were identified that are similar to GDSL lipases from a wide variety of plants, but which are not closely related to TcGLIP. The canSat3 scaffold encoding this enzyme (scaffold5401) appears to encode three different GSDL esterase/lipase homologs, but the assembly of that region is somewhat fragmentary. GDSL esterase/lipase are multifunctional enzymes that include numerous related family members in sequenced plant genomes – with over 100 copies in some genomes.^14^ It is unclear, therefore, whether the GDSL lipase we identified would catalyze the same reaction as TcGLIP. The catalytic Ser-Asp-His triad and two other amino acids that Kikuta et. al. identified as necessary for TcGLIP functionality are all preserved in the *Cannabis* GDSL homologue (figure 5).^13^

**Figure 5.**
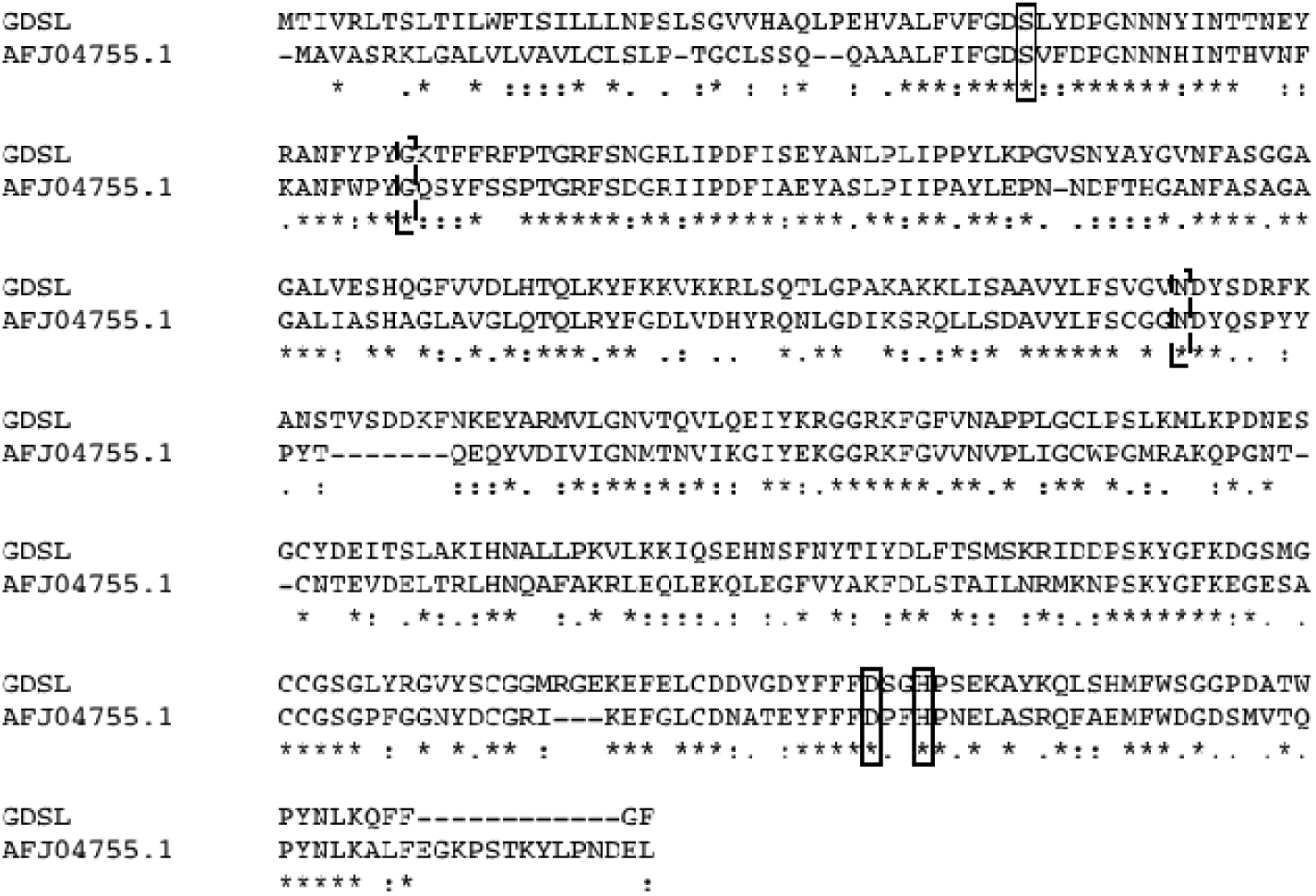
Clustal alignment of *Cannabis* GDSL lipase to the *T. cinerariifolium* GDSL lipase. The amino acids of catalytic triad that performs the esterification are boxed. Two other amino acids identified as significant for TcGLIP’s production of pyrethrins are indicated by a dashed box.^13^

## Discussion

Pyrethrins were found in cannabis flower from a sample grown in a closed environment, not exposed to exogenous pesticides. The transition from m/z = 329.3 to 161.1 is used to quantify the amount of pyrethrin I present, while a second transition from m/z = 329.3 to 133.2 allows the purity of the sample to be verified. The first transition will select for pyrethrin I, eliminating the other five forms of pyrethins. With respect to pyrethrin I, the ratio of these two transitions should be roughly 3:1. When cannabis flower samples are tested, however, ratios of 1:10 or 1:20 are common^∗^, meaning that some compound in cannabis has a retention time similar to pyrethrins, a molecular weight of 328-330 g/mol, and fragments to a 133 g/mol ion. Neither tetrahydrocannabinol (weight 314.5 g/mol) nor tetrahydrocannabinolic acid (weight 358.5 g/mol) fit this description. One possibility is that cannabis produces an isomer of pyrethrin I that more readily fragments to a 133 g/mol ion.

A number of enzymes are involved in producing pyrethrins in *T. cinerariifolium*, and many have been characterized genetically. Mapping the genes of these enzymes onto the *Cannabis* genome provides a plausible pathway by which cannabis could produce pyrethrins. One major issue is the homology of *Cannabis* transcripts with FDS rather than CDS. Whereas CDS causes cyclopropanation of two DMAPP units, FDS generally elongates a DMAPP moiety by adding an IPP unit.^9,15^ CDS appears unable to incorporate IPP onto the DMAPP moiety of a chain, although the mechanisms of CDS and FDS are very similar. Both FDS and CDS contain DDXXD motifs that coordinate Mg^2+^ ions to stabilize the substrates in the active site. A key feature of CDS is that the second of these motifs contains an asparagine residue instead of aspartate.^9^ This mutation is also present in the *Cannabis* FDS gene (albeit at a slightly different location), suggesting similarity to CDS (figure 4).

The other major step in the production of pyrethrins is the final step, in which a GDSL lipase-like enzyme forms the ester bond of pyrethrins. Although the alignment of the *Cannabis* homologue to TcGLIP was fragmentary, the homologue contained five key residues that are integral to pyrethrin production (figure 5). These include the Ser-Asp-His triad that forms the ester bond, as well as two amino acids that are necessary for pyrethrin production.^13^

These data are preliminary and the possibility that cannabis produces pyrethrins endogenously needs to be replicated by future studies. Such work could include expression of the genes identified herein, as well as larger controlled grows. Also missing is an explanation of the fragment with m/z = 133 that is identified at 30-60 fold high concentrations in cannabis flower than the pyrethrin standard. If cannabis indeed produces pyrethrins endogenously, this raises a number of significant medical and regulatory questions: how common are pyrethrin-producing cannabis plants? How do cultivation practices influence pyrethrin production? How should pyrethrins be regulated as a pesticide used on cannabis?

The last question is of particular significance to states legalizing medical and recreational cannabis. A number of states limit pyrethrins in cannabis to 1 μg/g or lower, whereas 2.48 μg/g was obtained from a “clean” plant in this work. The endogenous production of pyrethrins by cannabis would not necessarily imply that this concentration of pyrethrins is safe to smoke, vaporize, or ingest. But such data demonstrates the importance of a comprehensive approach to understanding the impact to consumers of pesticides in cannabis products.

## Footnotes

∗ According to conversations with Sonoma Lab Works.

